# Effect of topical soluble epoxide hydrolase inhibition in a preclinical diabetic wound healing model

**DOI:** 10.1101/571984

**Authors:** William C. Reisdorf, Neetu Rajpal, Andrew J. Gehman, Piyush Jain, Mark E. Burgert, Sujatha Sonti, Pankaj Agarwal, Deepak K. Rajpal

## Abstract

The objective of this study was to test a topical formulation of *EPHX2* inhibitor, GSK2256294, in a dermal wound diabetic *(db/db)* mouse model. Comparisons were made between groups treated with *EPHX2* inhibitor, matching vehicle control and a currently approved treatment for diabetic ulcers, becaplermin/Regranex, as a positive control which is highly efficacious in this model. Leptin receptor-deficient *(db/db)* male mice were sequentially assigned to treatment groups (N=10 per group) based on blood glucose levels and body weight. Mice were given a single 8 mm diameter full thickness wound on the back. Wounds were photographed and traced, and fresh test materials applied periodically with fresh dressing. Of ten mice treated with GSK2256294, three had completely healed wounds at the study conclusion, and 8/10 mice reached at least 50% closure. In the vehicle group, no mice achieved complete closure, and only 6/10 reached at least 50% closure by the study conclusion. Nine of ten wounds achieved complete closure in the positive control group by 21 days. Although the EPHX2 inhibitor results were indicative of efficacy, the single-sided p-value criterion of 0.05 was not met in this study.

## Introduction

Soluble epoxide hydrolase (sEH, encoded by the *EPHX2* gene), a member of the *α*/*β*-hydrolase fold family of epoxide hydrolases, is a critical enzyme in the metabolism of epoxyeicosatrienoic acids (EETs) to dihydroxyeicosatrienoic acids (DHETs). EETs are formed by the oxidation of arachidonic acid by CYP450 enzymes, primarily members of the CYP 2C and 2J classes in humans. EETs are a group of eight compounds: the four regioisomers are 5,6-EET, 8,9-EET, 11,12-EET and 14,15-EET, with R/S and S/R enantiomers of each. *In vitro*, EETs have demonstrated anti-inflammatory and potential pro-resolving effects as well as effects on airway and vascular smooth muscle, and vascular endothelial cells. The actions shown by EETs in animal models and human cellular assays include vasodilatory effects, ion channel modulation, mitogenesis and modulation of endothelial cell survival and vascular smooth muscle cell proliferation.

Although the molecular mechanisms of action of EETs have not been completely elucidated, it is hypothesized that the therapeutic effects of sEH inhibition are mediated by the accumulation of EETs and that many potentially beneficial actions of EETs are attenuated by the conversion of EETs to the less active DHETs. Therefore, sEH inhibition is a potential therapeutic target to enhance the beneficial effects of EETs (1). GSK2256294 is a potent and selective soluble epoxide hydrolase (sEH) inhibitor (2) which has previously been studied in preclinical models (3, 4) and in phase I clinical trials (5, 6).

Chronic wounds, including pressure ulcers, leg ulcers and diabetic foot ulcers are an area of significant unmet need. Thousands of lower limb amputations are performed annually on diabetic patients. One estimate of the direct cost of diabetes treatments in the US in 2007 was $116 billion, of which one third was linked to treating foot ulcers (7). Recognizing that diabetic wound healing is a very challenging paradigm, we hypothesized that inhibition of sEH would result in improved resolution of chronic wounds and could lead to a potentially transformational medicine for patients. We therefore proceeded to test GSK2256294 in a validated diabetic wound healing model to investigate this hypothesis.

Although there have been no GWAS focused on chronic wound healing, there are some interesting Mendelian skin disorders involving other enzymes of arachidonic acid metabolism. Autosomal recessive congenital ichthyosis type 1 (ARCI1, OMIM #242300) is caused by loss of function (LoF) mutation in *TGM1* (transglutaminase 1), ARCI2 (OMIM #242100) is caused by LoF in *ALOX12B* (arachidonate 12-lipoxygenase, 12R type) and ARCI3 (OMIM #606545) is caused by LoF in *ALOXE3* (arachidonate lipoxygenase 3). All three are keratinization disorders, characterized by abnormal skin scaling over the whole body (8). Mice with knockout expression of *TGM1, ALOX12B* or *ALOXE3* have impaired skin barrier function (9–11). EETs have been shown to activate *TGM1* in human and mouse keratinocytes (12), and *TGM1* mRNA facilitated wound healing in neonatal mouse skin (13).

Several published reports implicate direct involvement of sEH in wound healing in non-diabetic models. A tool sEH inhibitor (t-AUCB, *trans*-4-[4-(3-adamantan-1-yl-ureido)-cyclohexyloxy]-benzoic acid) significantly accelerated wound closure, epithelialization and neovascularization in an ear wound model in SKH-1 hairless mice (14), as did administration of sEH substrates 11,12-EET and 14,15-EET. The same research group showed accelerated wound closure in *EPHX2* KO mice (15). An independent study using knockout mice, sEH substrates and a different tool inhibitor (TUPS, 1-(1-methanesulfonyl-piperidin-4-yl)-3-(4-trifluoromethoxy-phenyl)-urea) has also shown similarly enhanced wound healing (16). In addition, it was demonstrated that increased expression of *CYP2C8* or *CYP2J2* also increased EET levels and promoted wound healing, whereas increased expression of sEH itself suppressed wound healing.

### Materials and methods

#### Animal Care

We contracted with Explora BioLabs, San Diego, CA to manage the animal study. Every effort was made to minimize or eliminate pain and suffering of all the animals in the study. The study was approved by the Office of Animal Welfare, Ethics and Strategy at GSK (Reference number D012050, Submission number S003940). All studies were conducted in accordance with the GSK Policy on the Care, Welfare and Treatment of Laboratory Animals and were reviewed by the Institutional Animal Care and Use Committee at Explora BioLabs. Male mice, *db/db* (Homozygous for Lepr^db^ <BKS.Cg-Dock7^m^ +/+ Lepr^db^/J>, 8-9 weeks old, were purchased from Jackson Laboratories. Upon arrival, animals were inspected for correct age and sex, ear-tagged and their body weights were recorded. The animals were housed 1-per cage. The animal housing, care and husbandry were performed per the Guide for the Care and Use of Laboratory Animals; 8th Edition; Institute of Laboratory Animal Resources; U.S. Department of Health and Human Services; National Institutes of Health Publication No. 85-23, Revised 2011.

Temperature in the animal holding room was maintained between 20 and 26°C (68-79°F). Room humidity was monitored (40-70% is recommended in the Guide). Temperature and humidity values were recorded daily. A 12-hour light/12-hour dark cycle illumination period was maintained. Animals were housed using the dual HEPA-filtered ventilated Innovive disposable IVC cage system 3.0, which provides 60 fresh-air changes per hour for animal cages. The animals had ad libitum access to standard diet (2920X.10, Global 18% Protein Rodent Diet from Harlan, San Diego, CA) and acidified water (pH 2.7-3.0) throughout the study period. The bedding material was hardwood chips (Sani-Chips, Cat # 7115, Harlan, CA, USA) and was changed weekly to ensure the ammonia levels within each cage were below 24 ppm.

Animals were acclimated to the housing environment for 7 days prior to the initiation of the study. During the acclimation period, the general health of the animals was monitored daily. Animals appeared normal and did not exhibit signs of poor health. No animals were excluded from this study. Glucose and body weight were measured on Day -1, and treatment groups were assigned sequentially to animals ordered by their Day -1 glucose levels (e.g., 1, 2, 3, 4, 5, 5, 4, 3, 2, 1, etc.) to obtain similar average blood glucose and body weight values among groups. The treatment group assignment was performed on Study Day -1 prior to test article administration. Animals were weighed once per week, and body weights were recorded. The general health condition and attitude of each animal was monitored daily throughout the study. Clinical signs indicative of poor health, stress and pain were noted.

#### Topical sEH Formulation

Topical gel formulations for GSK2256294 (2) were developed by Dermatology. The solvent system was decided based on solubility of GSK2256294. Gel formulation prototypes were then prepared with the solvent system using various combinations of polymers and pH adjusters.

The formulation used in the animal study comprised of 1% (%w/w) GSK2256294, 20% PEG400, 25% propylene glycol, 25% transcutol, 1% Ultrez-10, 1.5% triethanolamine in water. Vehicle control lacked GSK2256294 (q.s. with water) but was otherwise identical. Fresh supply of the topical formulation was shipped to the CRO weekly, for the duration of the study. Becaplermin (0.01%) was purchased from Smith & Nephew (Catalogue number - REG810-15) and used as directed.

#### Study Design

We requested a representative data set from the CRO to assess model variability and estimate the number of mice necessary to achieve adequate power. Sample sizes of N=10 mice per group were determined to achieve 80% power to detect decreases from vehicle control of 1.31 days in mean interpolated time to 50% wound closure when using a one-way ANOVA model with five experimental groups, for a total sample size of 50 animals. The study contained five experimental groups in order to investigate other chemical compounds in addition to GSK2256294, although only the three previously mentioned groups are presented here.

#### Timeline

- Day -7 to Day 0 acclimation
- Day -3 back hair removed
- Day 1 wounding
- Day 1 pre-dose and 1 hour post-dose blood collection (N=3 mice per group)
- Days 1, 4, 7, 10, 13, 16, 19, 21 and 24 dosing and wound measurement
- Day 21 termination of positive control group
- Day 26 termination of sEHi and vehicle test groups, wound measurement

On Day -3, all animals were anesthetized via inhaled isoflurane and the entire back hair shaved with clippers. The remaining hair was removed with depilatory cream (Nair, unscented/sensitive skin). On Day 1, animals were anesthetized via inhaled isoflurane and a single 8mm diameter full thickness (skin and the underlying panniculus carnosus) wound created starting 1cm below the scapula bones running parallel at 0.5cm lateral from the spine. Animals received ~50-100μL of Positive Control or Test Article or Vehicle spread out to create an even and consistent layer over the wound making sure to cover all the wound area. Wounds were covered with an occlusive dressing [High MVP Transparent Dressing (Smith + Newman) and Mastisol liquid adhesive (Eloquent Healthcare)] and changed on dosing days following surgery. Animals were weighed once per week, and body weights recorded. Weight loss in excess of 20% compared to Day 0 was to be reported, but no mice reached this threshold during the study. On Study Day 21, the animals in the positive control group were euthanized by isoflurane overdose. On Study Day 26, the animals in the sEHi GSK2256294 treatment and vehicle control groups were euthanized. Wounds were collected into 10% formalin for storage. Each wound was excised completely to include underneath muscle and 5mm of surrounding tissue. The personnel conducting the animal wounding, dosing, and measuring of wound area were blinded to the identity of each test article, although the laboratory was aware of this information as part of study planning and reporting. The dosing, photographing, and measuring of wound area were each conducted sequentially by group and not in a random order.

#### Data Analysis

Wounds were photographed, traced and analyzed using the computer program Image J software from digital photographs (www.rsb.info.nih.gov/ij/). Analysis was performed using R package *lme4* version 1.1-18-1 and SAS/STAT version 12.1. The tests of statistical significance are reported for a single sided test in the direction of reduced wound size or time to smaller wound size. All endpoints for statistical analysis were derived from the wound size measurements and were based on individual mice being the experimental unit. Data from all five experimental groups were included in these analyses. A variety of alternative statistical approaches were considered instead of the one-way ANOVA used for the earlier power analysis due to some mice not achieving 50% closure during the study.

Time to 50% closure (T_50_) was estimated using a spline model to estimate the wound closure percent at timepoints between the days of measurement. The spline model incorporated a forced intercept of 100% at Day 1 to account for the normalization of wound area to the baseline measurement. It also involved knots at Days 4, 10, 16 and 21, a term for the interaction between group and the spline, and a random coefficient for Day that varied by animal as repeated measurements were observed for each mouse. Group means and 90% confidence intervals were computed. For each group, the T_50_ mean estimate and its confidence intervals were calculated from the intersection of the spline model estimates with 50% closure. Time to 50% closure was also evaluated using survival analysis due to some mice not achieving 50% closure during the study. The time of event of 50% closure was taken as the observed Study Day when a wound was at or below half its baseline size. A separate analysis calculated the time of event using linear interpolation connecting Study Days between which 50% closure was achieved. Wounds which remained larger than 50% of their baseline on all observed Study Days were considered censored (no event). Time to complete closure was similarly analyzed using the first observed Study Day when the wound was completely closed as the time to event. The log-rank test was used to compare treatment groups by their survival.

Baseline-normalized wound area under the curve (AUC) was also computed for each mouse, and group means and standard errors were reported, using a one-way ANOVA. Similarly, wound area AUC was calculated without normalization and was modeled by an ANCOVA model, with baseline wound area as the covariate.

### Results

Results for time to 50% closure using the spline model are shown in Figure 1. The positive control 0.01% becaplermin (Regranex) group showed complete healing in 9/10 animals over the course of 21 days. The T_50_ (the time to 50% closure estimate) was 11.5 days, with 90% confidence limits of approximately 11 and 12 days. The positive control arm was terminated at Day 21. The GSK2256294 treatment group showed healing, but the process was slower than for the positive control group and not all animals responded: T_50_ = 14.4 days, 90% confidence limits of 13 and 16.5 days. The vehicle group also showed some healing over time, T_50_ = 19.7 Days, 90% confidence lower time 15.5 days (insufficient data to estimate the upper confidence limit), but the progress was somewhat slower than the GSK2256294 group.

**Fig 1.**
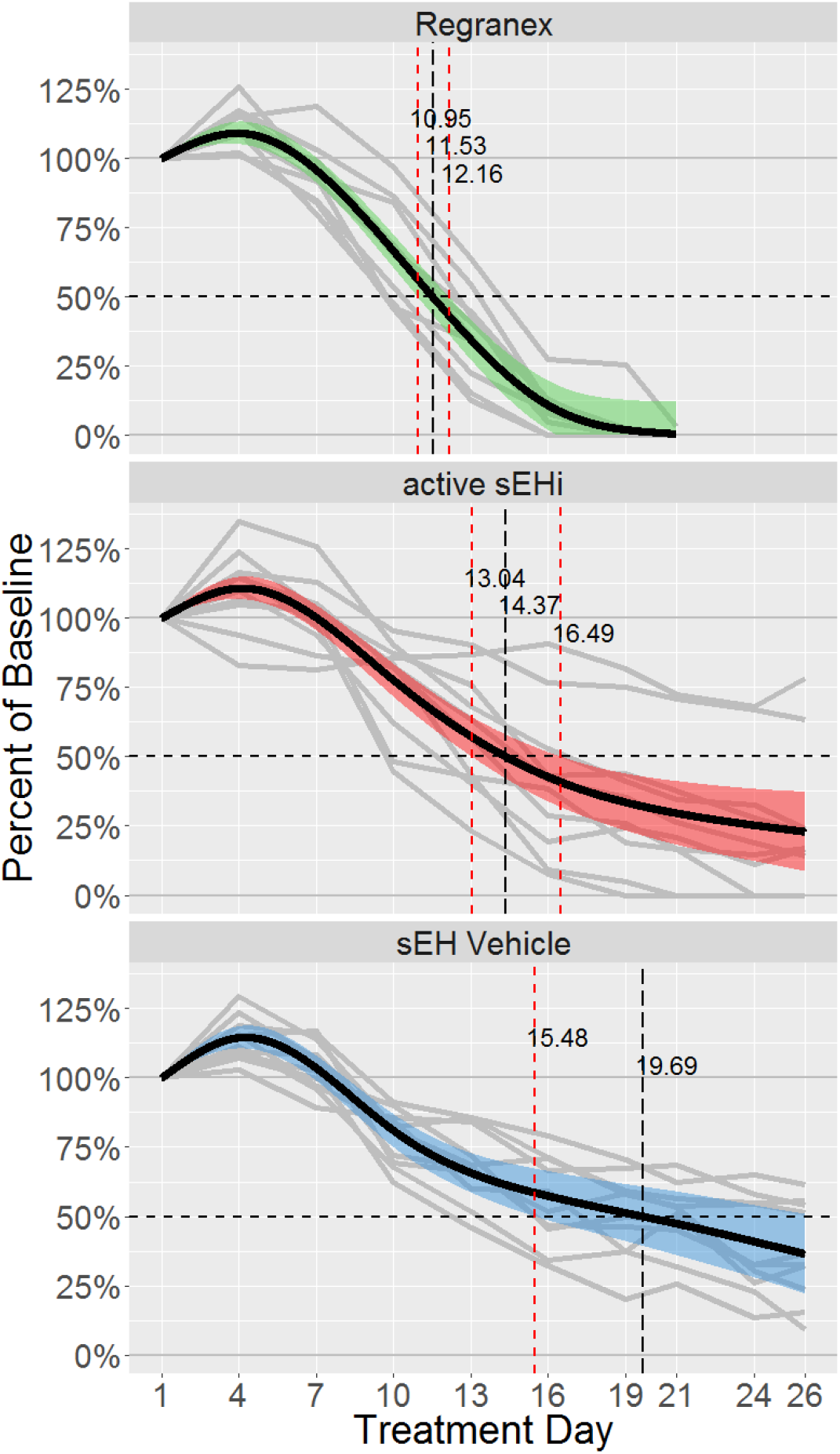
Time to 50% wound closure (T_50_) for topical GSK2256294. *db/db* mice (N=10 per treatment group) were treated with topical GSK2256294, vehicle control, or positive control (0.01% becaplermin). Wound areas were determined by tracing the wound boundary at each time point and compared to the wound area on Day 1 (baseline = 100%). A spline model was used to interpolate between the discrete wound area measurements. The dashed horizontal line indicates 50% closure. The dashed black vertical line shows the treatment group estimate of mean T_50_, and the red dashed vertical lines show the 90% confidence interval for T_50_. Grey lines represent data from individual animals.

Figure 2 shows the individual values and group means for baseline-normalized wound AUC through Day 21, by which time the positive control group wounds had healed. The mean AUC of the positive control group (shown in green) was significantly different (11.1 cm^2^ total AUC [90% confidence interval: 9.6-12.5] vs. 13.3 [12.8-15.8]; p-value = 0.002) than the GSK2256294 group (shown in red). However, the GSK2256294 group was not significantly different from the vehicle control group (shown in blue; 13.3 [12.8-15.8] vs. 15.9 [14.5-17.4]; p-value = 0.061). The same conclusion was reached when wound AUC through Day 21 was modeled with baseline as the covariate (not shown): the mean AUC of the GSK2256294 treated group was significantly higher than that of the positive control group (9.3 [8.4-10.2] vs. 7.2 [6.3-8.1]; p-value = 0.001) but not significantly lower than the vehicle control group (9.3 [8.4-10.2] vs. 10.3 [9.3-11.2], respectively; p-value = 0.075).

**Fig 2.**
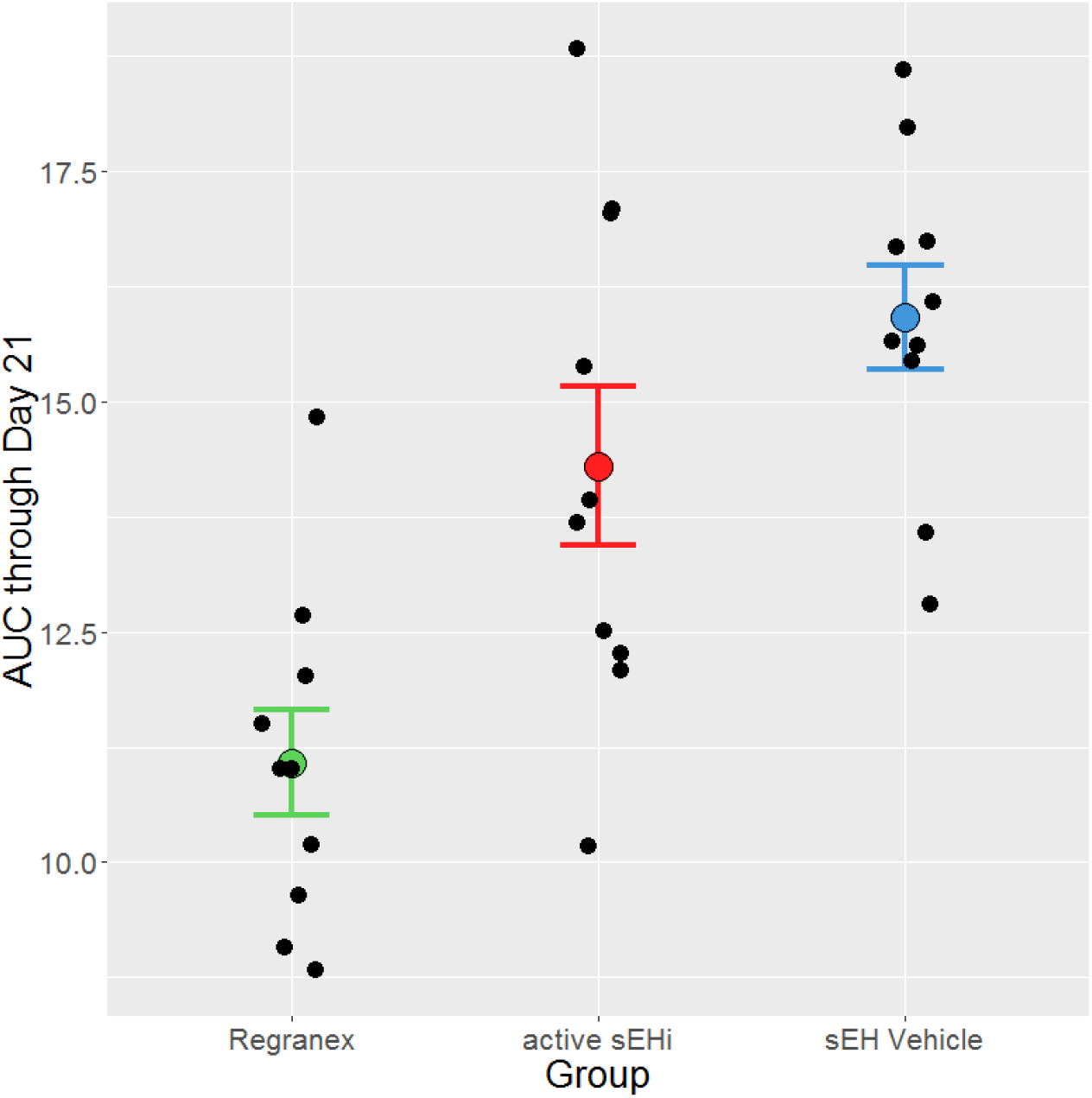
Wound AUC through day 21 for topical GSK2256294. Data points show the computed AUC of wound area self-normalized to Day 1 baseline, for individual mice in the sEHi (red), vehicle (blue) and positive control (green) groups. Group means and standard error bars are also displayed.

Because healing appeared to still be progressing at Day 21, the study was extended until Day 26 for the sEHi and vehicle arms, to determine if the healing trend would continue. The Day 26 results are shown in Figure 3. While the GSK2256294-treated group appeared to show more healing than the vehicle control, the results did not meet the criterion for statistical significance (15.6 [13.4-17.8] vs. 18.0 [15.8-20.2]; p-value = 0.058). Similarly, in comparison of wound AUC through Day 26 with baseline area as the covariate (not shown), the mean AUC of the GSK2256294 treated group was not significantly lower than that of the vehicle control group (10.0 [8.6-11.5] vs. 11.6 [10.1-13.0]; p-value = 0.067).

**Fig 3.**
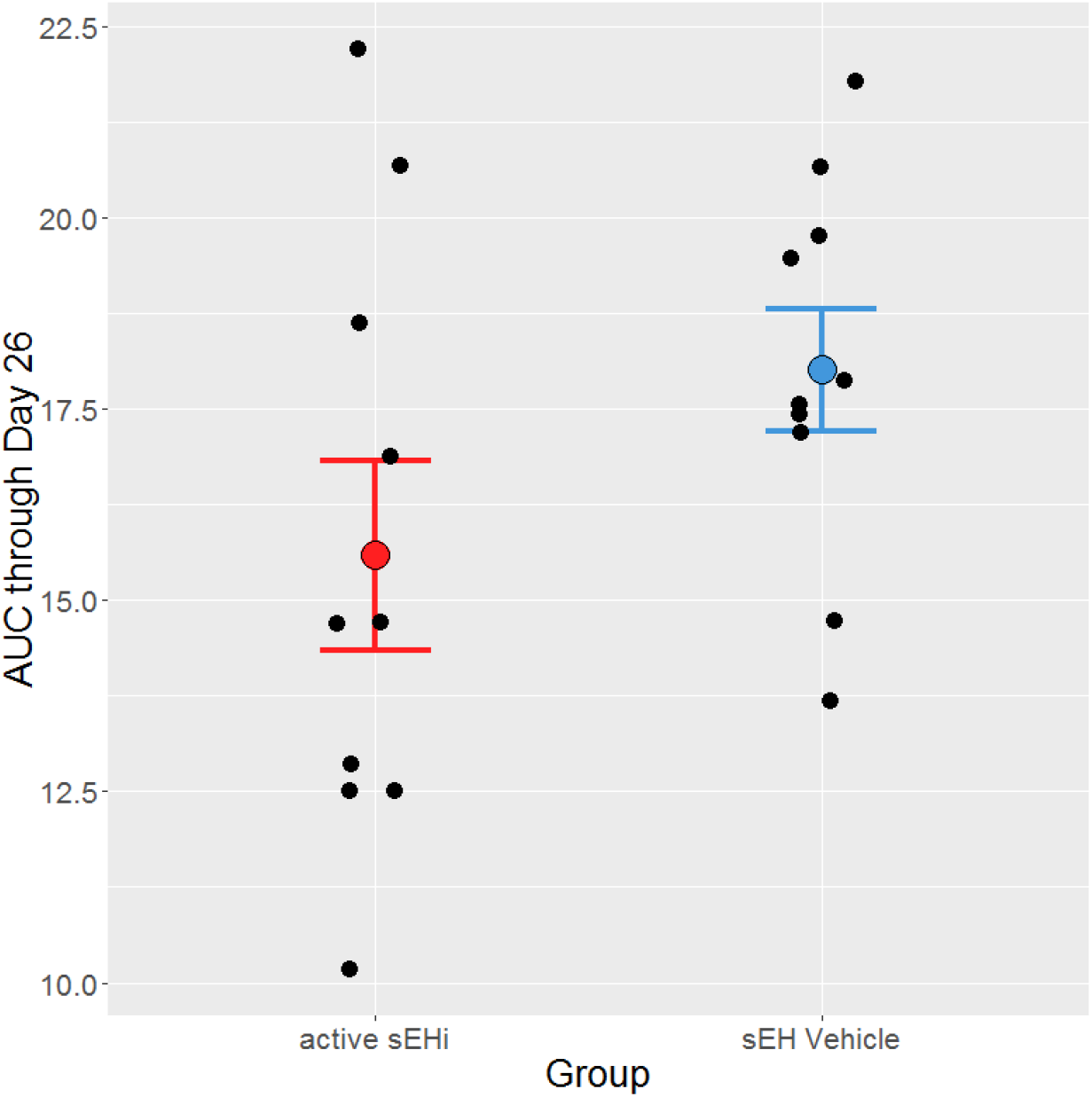
Wound AUC through day 26 for topical GSK2256294. Data points show the computed AUC of wound area self-normalized to Day 1 baseline, for individual mice in the sEHi (red) and vehicle (blue) groups. Group means and standard error bars are also displayed.

To summarize, 9/10 mice in the positive control group achieved wound closure by day 21, 3/10 mice in the GSK2256294 treatment group achieved closure by Day 26 and no mice in the vehicle group achieved closure by Day 26 (Table 1). The wound area data are available in Supplemental Table 1.

**Table 1.** Summary of results for wound closure

| **Day 21** |  |  |  |
| --- | --- | --- | --- |
| **Treatment Group** | **% wound closure (median)** | **N 100% closure** | **N 50% closure** |
| **becaplermin (positive control)** | 100 | 9/10 | 10/10 |
| **Active sEHi** | 76 | 2/10 | 8/10 |
| **Vehicle control** | 48 | 0/10 | 6/10 |

| **Day 26** |  |  |  |
| --- | --- | --- | --- |
| **Treatment Group** | **% wound closure (median)** | **N 100% closure** | **N 50% closure** |
| **Active sEHi** | 83 | 3/10 | 8/10 |
| **Vehicle control** | 65 | 0/10 | 6/10 |

## Discussion

We were encouraged that the GSK2256294-treated group showed evidence of improved healing compared to vehicle control, despite the statistical significance threshold of P < 0.05 not being met. In part, this might be attributed to greater variability in the wound healing process among the mice in this study, compared to baseline data previously seen in this model. Repeating the study with larger group size would be illuminating. We also note that this study was our first attempt at topical formulation of GSK2256294. It may be possible to further optimize the formulation, and the dose administered, to obtain improved response.

After our study was complete, a publication on EETs and diabetic wound healing appeared (17), demonstrating that 11,12-EET treatment accelerated wound healing in *ob/ob* (leptin-deficient) mice. These mice also had significantly reduced mRNA and protein expression of *CYP2C65* and *CYP2J2* compared to wild type. Taken together, the data strongly suggests that sEH inhibition, or other strategies to raise EET levels, can potentially improve wound healing in pre-clinical diabetic models.

The positive control Becaplermin is a biological agent (recombinant human platelet-derived growth factor-BB), which has been approved and launched for use in treating diabetic ulcers. Although it works well in the *db/db* mouse model, its efficacy in patients is less impressive. A meta-analysis of five clinical trials (18) found that 205/428 (47.89%) of becaplermin-treated ulcers healed completely, versus 109/335 (32.53%) of those in the placebo group. Considering that becaplermin has a black box warning for malignancies in patients using more than three tubes, there is clearly unmet need in the treatment of diabetic ulcers. Finally, we point out that there is additional evidence from non-diabetic mouse models that pre-clinical sEH inhibitors improve wound healing in that context (14–16), so applications for this therapy could be extended to surgical and other types of wounds.

## Acknowledgements

2017 The authors wish to acknowledge our colleagues for their support and critical discussions: Spiro Getsios, Laure Rittié, Akanksha Gupta and Jennifer Singh. We also thank Yolanda Sanchez for making available EPHX2 inhibitor GSK2256294 for testing in wound healing, and for insightful comments on earlier drafts of this manuscript.

## Supporting Information

S1 Table. Wound healing time-course measurements for individual mice in each treatment group.

