## Supplemental Table 1 for "Effect of topical soluble epoxide hydrolase inhibition in a preclinical diabetic wound healing model"

| | Date: | 1/24/17 | | 1/27/17 | | 1/30/17 | | 2/2/17 | | 2/5/17 | | 2/8/17 | | 2/11/17 | | 2/13/17 | | 2/16/17 | | 2/18/17 | |
|---|---|---|---|---|---|---|---|---|---|---|---|---|---|---|---|---|---|---|---|---|---|
| | Study Day: | 1 | | 4 | | 7 | | 10 | | 13 | | 16 | | 19 | | 21 | | 24 | | 26 | |
| **Group** | **Animal ID** | **Wound Area ($cm^2$)** | **% Wound Closure** | **Wound Area ($cm^2$)** | **% Wound Closure** | **Wound Area ($cm^2$)** | **% Wound Closure** | **Wound Area ($cm^2$)** | **% Wound Closure** | **Wound Area ($cm^2$)** | **% Wound Closure** | **Wound Area ($cm^2$)** | **% Wound Closure** | **Wound Area ($cm^2$)** | **% Wound Closure** | **Wound Area ($cm^2$)** | **% Wound Closure** | **Wound Area ($cm^2$)** | **% Wound Closure** | **Wound Area ($cm^2$)** | **% Wound Closure** |
| | 415 | 0.612 | 0 | 0.662 | -8 | 0.627 | -2 | 0.382 | 38 | 0.281 | 54 | 0.198 | 68 | 0.123 | 80 | 0.158 | 74 | 0.083 | 86 | 0.094 | 85 |
| | 430 | 0.529 | 0 | 0.685 | -29 | 0.603 | -14 | 0.435 | 18 | 0.454 | 14 | 0.418 | 21 | 0.372 | 30 | 0.329 | 38 | 0.344 | 35 | 0.326 | 38 |
| | 445 | 0.651 | 0 | 0.725 | -11 | 0.624 | 4 | 0.441 | 32 | 0.336 | 48 | 0.224 | 66 | 0.242 | 63 | 0.208 | 68 | 0.149 | 77 | 0.061 | 91 |
| | 470 | 0.609 | 0 | 0.752 | -23 | 0.650 | -7 | 0.438 | 28 | 0.412 | 32 | 0.285 | 53 | 0.283 | 54 | 0.274 | 55 | 0.198 | 67 | 0.200 | 67 |
| 1 | 450 | 0.606 | 0 | 0.664 | -10 | 0.656 | -8 | 0.490 | 19 | 0.415 | 32 | 0.429 | 29 | 0.358 | 41 | 0.343 | 43 | 0.333 | 45 | 0.337 | 44 |
| sEHi | 458 | 0.594 | 0 | 0.686 | -15 | 0.694 | -17 | 0.412 | 31 | 0.393 | 34 | 0.307 | 48 | 0.347 | 42 | 0.316 | 47 | 0.179 | 70 | 0.141 | 76 |
| | 424 | 0.615 | 0 | 0.711 | -16 | 0.597 | 3 | 0.520 | 15 | 0.370 | 40 | 0.364 | 41 | 0.229 | 63 | 0.314 | 49 | 0.160 | 74 | 0.199 | 68 |
| | 413 | 0.779 | 0 | 0.803 | -3 | 0.695 | 11 | 0.667 | 14 | 0.655 | 16 | 0.520 | 33 | 0.526 | 32 | 0.534 | 31 | 0.452 | 42 | 0.421 | 46 |
| | 435 | 0.720 | 0 | 0.773 | -7 | 0.717 | 0 | 0.647 | 10 | 0.518 | 28 | 0.327 | 55 | 0.358 | 50 | 0.349 | 52 | 0.235 | 67 | 0.264 | 63 |
| | 444 | 0.653 | 0 | 0.776 | -19 | 0.756 | -16 | 0.595 | 9 | 0.561 | 14 | 0.465 | 29 | 0.380 | 42 | 0.351 | 46 | 0.355 | 46 | 0.336 | 49 |
| | 422 | 0.573 | 0 | 0.671 | -17 | 0.592 | -3 | 0.496 | 13 | 0.311 | 46 | 0.066 | 88 | 0.000 | 100 | 0.000 | 100 | - | - | - | - |
| | 464 | 0.671 | 0 | 0.736 | -10 | 0.535 | 20 | 0.321 | 52 | 0.102 | 85 | 0.000 | 100 | 0.000 | 100 | 0.000 | 100 | - | - | - | - |
| | 433 | 0.646 | 0 | 0.747 | -16 | 0.630 | 2 | 0.421 | 35 | 0.264 | 59 | 0.079 | 88 | 0.017 | 97 | 0.000 | 100 | - | - | - | - |
| | 440 | 0.662 | 0 | 0.832 | -26 | 0.619 | 6 | 0.556 | 16 | 0.284 | 57 | 0.031 | 95 | 0.000 | 100 | 0.000 | 100 | - | - | - | - |
| 2 | 419 | 0.625 | 0 | 0.717 | -15 | 0.743 | -19 | 0.606 | 3 | 0.398 | 36 | 0.172 | 72 | 0.160 | 74 | 0.020 | 97 | - | - | - | - |
| Regranex | 442 | 0.635 | 0 | 0.690 | -9 | 0.630 | 1 | 0.437 | 31 | 0.226 | 64 | 0.031 | 95 | 0.000 | 100 | 0.000 | 100 | - | - | - | - |
| | 436 | 0.748 | 0 | 0.759 | -1 | 0.635 | 15 | 0.402 | 46 | 0.168 | 78 | 0.067 | 91 | 0.000 | 100 | 0.000 | 100 | - | - | - | - |
| | 468 | 0.641 | 0 | 0.653 | -2 | 0.538 | 16 | 0.294 | 54 | 0.081 | 87 | 0.000 | 100 | 0.000 | 100 | 0.000 | 100 | - | - | - | - |
| | 448 | 0.681 | 0 | 0.739 | -9 | 0.631 | 7 | 0.315 | 54 | 0.234 | 66 | 0.054 | 92 | 0.000 | 100 | 0.000 | 100 | - | - | - | - |
| | 417 | 0.739 | 0 | 0.744 | -1 | 0.678 | 8 | 0.494 | 33 | 0.330 | 55 | 0.097 | 87 | 0.000 | 100 | 0.000 | 100 | - | - | - | - |
| | 463 | 0.630 | 0 | 0.522 | 17 | 0.512 | 19 | 0.540 | 14 | 0.548 | 13 | 0.571 | 9 | 0.514 | 18 | 0.457 | 27 | 0.430 | 32 | 0.494 | 22 |
| | 426 | 0.620 | 0 | 0.711 | -15 | 0.652 | -5 | 0.509 | 18 | 0.352 | 43 | 0.177 | 71 | 0.160 | 74 | 0.102 | 84 | 0.090 | 85 | 0.106 | 83 |
| | 434 | 0.634 | 0 | 0.855 | -35 | 0.797 | -26 | 0.574 | 9 | 0.433 | 32 | 0.336 | 47 | 0.261 | 59 | 0.219 | 65 | 0.206 | 68 | 0.151 | 76 |
| | 460 | 0.610 | 0 | 0.756 | -24 | 0.627 | -3 | 0.509 | 17 | 0.266 | 56 | 0.057 | 91 | 0.029 | 95 | 0.000 | 100 | 0.000 | 100 | 0.000 | 100 |
| 3 | 428 | 0.639 | 0 | 0.680 | -6 | 0.650 | -2 | 0.308 | 52 | 0.273 | 57 | 0.246 | 62 | 0.121 | 81 | 0.105 | 84 | 0.000 | 100 | 0.000 | 100 |
| sEH vehicle | 451 | 0.683 | 0 | 0.791 | -16 | 0.666 | 2 | 0.305 | 55 | 0.159 | 77 | 0.053 | 92 | 0.000 | 100 | 0.000 | 100 | 0.000 | 100 | 0.000 | 100 |
| | 429 | 0.769 | 0 | 0.721 | 6 | 0.666 | 13 | 0.635 | 17 | 0.487 | 37 | 0.319 | 59 | 0.272 | 65 | 0.203 | 74 | 0.147 | 81 | 0.110 | 86 |
| | 437 | 0.653 | 0 | 0.760 | -16 | 0.738 | -13 | 0.622 | 5 | 0.589 | 10 | 0.502 | 23 | 0.490 | 25 | 0.463 | 29 | 0.436 | 33 | 0.415 | 36 |
| | 469 | 0.632 | 0 | 0.662 | -5 | 0.650 | -3 | 0.551 | 13 | 0.479 | 24 | 0.272 | 57 | 0.277 | 56 | 0.237 | 63 | 0.177 | 72 | 0.145 | 77 |
| | 438 | 0.654 | 0 | 0.718 | -10 | 0.613 | 6 | 0.408 | 38 | 0.263 | 60 | 0.127 | 81 | 0.159 | 76 | 0.137 | 79 | 0.074 | 89 | 0.109 | 83 |
